# Modulating dream experience: Noninvasive brain stimulation over the sensorimotor cortex reduces dream movement

**DOI:** 10.1101/600288

**Authors:** Valdas Noreika, Jennifer M. Windt, Markus Kern, Katja Valli, Tiina Salonen, Riitta Parkkola, Antti Revonsuo, Ahmed A. Karim, Tonio Ball, Bigna Lenggenhager

## Abstract

Recently, cortical correlates of specific dream contents have been reported, such as the activation of the sensorimotor cortex during dreamed hand clenching. Yet, the causal mechanisms underlying specific dream content remain largely elusive. Here, we investigated how alterations in the excitability of sensorimotor areas through transcranial direct current stimulation (tDCS) might alter dream content. Following bihemispheric tDCS or sham stimulation, participants who were awakened from REM sleep filled out a questionnaire on bodily sensations in dreams. tDCS, compared to sham stimulation, significantly decreased reports of dream movement, especially repetitive actions. Contrary to this, other types of bodily experiences, such as tactile or vestibular sensations, were not affected by tDCS, confirming the specificity of stimulation effects. In addition, tDCS reduced interhemispheric coherence in parietal areas and altered the phasic electromyography correlation between the two arms. These findings reveal that a complex reorganization of the motor network co-occurred with the reduction of dream movement, confirming spatial specificity of the stimulation site. We conclude that tDCS over the sensorimotor cortex causally interferes with dream movement during REM sleep.

## Introduction

Dreams are vivid, often emotionally intense and narratively complex experiences occurring in sleep. In our dreams, we feel immersed in alternative worlds and have the experience of interacting with other persons and objects. Often this involves the subjective experience of moving through the dream world, and movement is among the most frequently reported dream experiences, second only to visual imagery. Yet these rich subjective experiences stand in stark contrast to the outward unresponsiveness and lack of observable behaviour during sleep. This study aimed to investigate the causal mechanisms underlying dream movement and bodily experience in dreams by using tDCS over sensorimotor areas. While most existing studies of the neural underpinnings of bodily experience in dreams and dream movement are correlational, our approach allowed us to manipulate dream content and draw conclusions about its underlying causes.

Specifically, our goal was to characterize the role of sensorimotor cortex in the generation of bodily sensations in dreams. We aimed to experimentally inhibit motor and other bodily experiences as an important aspect of self-simulation in dreams through bihemispheric transcranial direct current stimulation (tDCS) during REM sleep. After awakening from REM sleep, subjective dream experience was examined through the collection of dream reports and a questionnaire specifically designed to investigate bodily experiences in dreams; neural measures were obtained through electrophysiological sleep data.

This experimental protocol was guided by theoretical and empirical considerations. Our focus on bodily experience was motivated by the centrality of self-experience and subjective presence to dreaming (Strauch and Meier 1996; Occhionero et al. 2005; Speth et al. 2013). The immersive structure of dreaming is foregrounded in simulation theories (Revonsuo et al. 2015), in which dreams are described as mental simulations characterized by the experience of a virtual world. Typically, this virtual world is centered on a virtual self and experienced from an internal first-person perspective. The dream self is typically described as actively engaged in dream events, and movement is reported in 75% of dreams (Strauch and Meier 1996; Cicogna and Bosinelli 2001). This immersive *here and now* quality is regarded as a defining characteristic of dreaming. It is also striking that with few exceptions, both the virtual world and the virtual self in dreams are experienced as real. Simulation views advocate the idea that “being-in-a-dream” feels the same as “being-in-the-world” during wakefulness. Moreover, bodily experience and movement sensations appear to be central to the feeling of subjective presence both during the waking and dream state, and sensorimotor interaction modulates subjective presence both in real and virtual environments (Sanchez-Vives and Slater 2005).

Our focus on bodily experience was further guided by findings suggesting high-level activity of the motor cortex during REM sleep (Hobson 1988; Maquet et al. 2000; Dang-Vu et al. 2005). Generally, REM sleep dreaming has been associated with relative deactivation of executive networks and frontal areas, and with high levels of activity in sensory, motor, and emotional networks as compared to wakefulness (Schwartz and Maquet 2002; Nir and Tononi 2010; Cipolli et al. 2017). Studies focusing on the neural correlates of specific types of bodily dream experiences have shown the sensorimotor cortex to be activated during hand clenching in dreams (Dresler et al. 2011), and the right superior temporal sulcus, a region involved in the biological motion perception, to be activated in dreams with a sense of movement (Siclari et al. 2017). Furthermore, smooth pursuit eye movements during tracking of a visual target are highly similar during waking perception and lucid REM sleep dreaming (LaBerge et al. 2018). Taken together, these studies suggest a remarkable isomorphism of the neural mechanisms underlying motor control in wakefulness and dreaming. However, the correlative nature of these studies limits their potential to uncover the causal contribution of specific brain regions to dream content.

Older studies attempted to experimentally induce different kinds of dream experience via peripheral and bodily stimulation during sleep. Causal manipulations that have been shown to have an effect on dream content include vestibular stimulation in rotating chairs (Hoff 1929; Hoff and Pötzl 1937) or hammocks (Leslie and Ogilvie 1996); light flashes or sprays of water applied to the skin (Dement and Wolpert 1958); thermal stimulation (Baldridge et al. 1965; Baldridge 1966); tactile stimulation via a blood pressure cuff inflated on the leg (Nielsen 1993; Sauvageau et al. 1998); and olfactory stimulation (Schredl et al. 2009). The frequency of stimulus incorporation in dreams is variable and dependent both on the kind of stimulus and the sensory modality. Particularly high incorporation rates were achieved in studies using blood pressure cuff stimulation (40-80%) (Nielsen 1993; Sauvageau et al. 1998). This method of causally manipulating dream content is promising. However, because the processing of external and peripheral stimuli is attenuated in REM sleep, the precise effect of sensory stimulation on dream content is often nonspecific and unpredictable.

As a more direct method for manipulating dream content that avoids the possibly distorting effect of reduced sensory processing during REM sleep, we previously suggested using tDCS (Noreika et al. 2010). We argued that this method might complement previous attempts to manipulate dream content through sensory and bodily stimulation in sleep. Unihemispheric tDCS has been shown to facilitate motor imagery during REM sleep (Speth and Speth 2016) and to modulate visual imagery during Stage 2 NREM sleep (Jakobson, Fitzgerald, et al. 2012a), but not during slow wave sleep (Jakobson, Fitzgerald, et al. 2012b) or REM sleep (Jakobson, Conduit, et al. 2012). Furthermore, frontal tDCS increases lucidity in experienced lucid dreamers (Stumbrys et al. 2013); and frontal transcranial alternating current stimulation (tACS) increases dissociation, insight and control in novice lucid dreamers (Voss et al. 2014). tDCS has also been reported to modulate mind wandering in wakefulness (Axelrod et al. 2015). This is promising, as dreaming has been proposed to be an intensified form of mind wandering, based on phenomenological and neurophysiological similarities (Fox et al. 2013).

Here, we applied tDCS over the sensorimotor cortex, aiming to understand its causal role in dream content generation. Since tDCS modulates neural processes associated with motor imagery during wakefulness (Quartarone et al. 2004; Matsumoto et al. 2010; Feurra, M. et al. 2011), we expected a similar effect during REM sleep. However, instead of planned facilitation of movement sensations in dreams with unilateral anodal tDCS (Speth and Speth 2016), our stimulation protocol was designed to interfere with motor processing during sleep, enabling a more focused analysis of the electrophysiological mechanisms underlying dream movement. Given that unilateral cathodal tDCS does not disrupt motor imagery during REM sleep (Speth and Speth 2016), we adopted a bihemispheric tDCS protocol, which is known to interfere with cortical and cerebellar motor networks more effectively than unilateral tDCS, particularly when applied during the resting state (Lindenberg et al. 2013, 2016).

To investigate possible effects of bihemispheric tDCS on outward muscular activity, we obtained electromyographic (EMG) measures from the arms. REM sleep is typically characterized by near-complete muscle atonia (Pompeiano 1967) and a partial blockade of sensory input (Hobson 1988; Wu 1993). At the same time, subtle muscular activity in the form of twitching is frequent in REM sleep and may play a role in the development and maintenance of motor behaviour (Blumberg 2015). A relation to dreaming seems plausible, but remains incompletely understood (Windt 2018).

We hypothesized that if the sensorimotor cortex has a causal role in generating sensorimotor dream content, bihemispheric tDCS over the sensorimotor cortex during REM sleep should attenuate movement and other bodily experiences in dreams reported immediately after timed awakenings in the laboratory. To test this hypothesis, we developed an empirically informed questionnaire focused specifically on bodily sensations in dreams. This allowed us to probe bodily experiences more systematically than the more common methods of content analysis or quantitative linguistic analysis of dream reports (Speth and Speth 2016). Furthermore, we hypothesized that bihemispheric tDCS during REM sleep would interfere with interhemispheric motor networks as well as with spontaneous peripheral muscle activity, which are possible neural pathways to the reduction of dream movement.

## Methods and Materials

### Methods outline

The study protocol consisted of a recruitment and screening session, an MRI session, and two sleep sessions on non-consecutive nights (see Figure 1A). In addition, a TMS assessment of motor cortical excitability took place on the evening of the first sleep session. Ten participants were awakened from REM sleep two or three times per night and asked to give free dream reports and to answer to the Bodily Experiences in Dreams (BED) Questionnaire, which targeted the dream immediately preceding awakening (see Figure 1B). Participants received sham-stimulation during REM sleep on one night and bihemispheric tDCS on the other night. Bihemispheric tDCS montage included a cathode placed over the left sensorimotor cortex and an anode placed over the right sensorimotor cortex (see Figure 1C). In addition to standard polysomnography, central and peripheral electrophysiological data were recorded using 16 EEG channels and 4 EMG channels measuring flexor and deltoid muscles in both arms.

**Figure 1.**
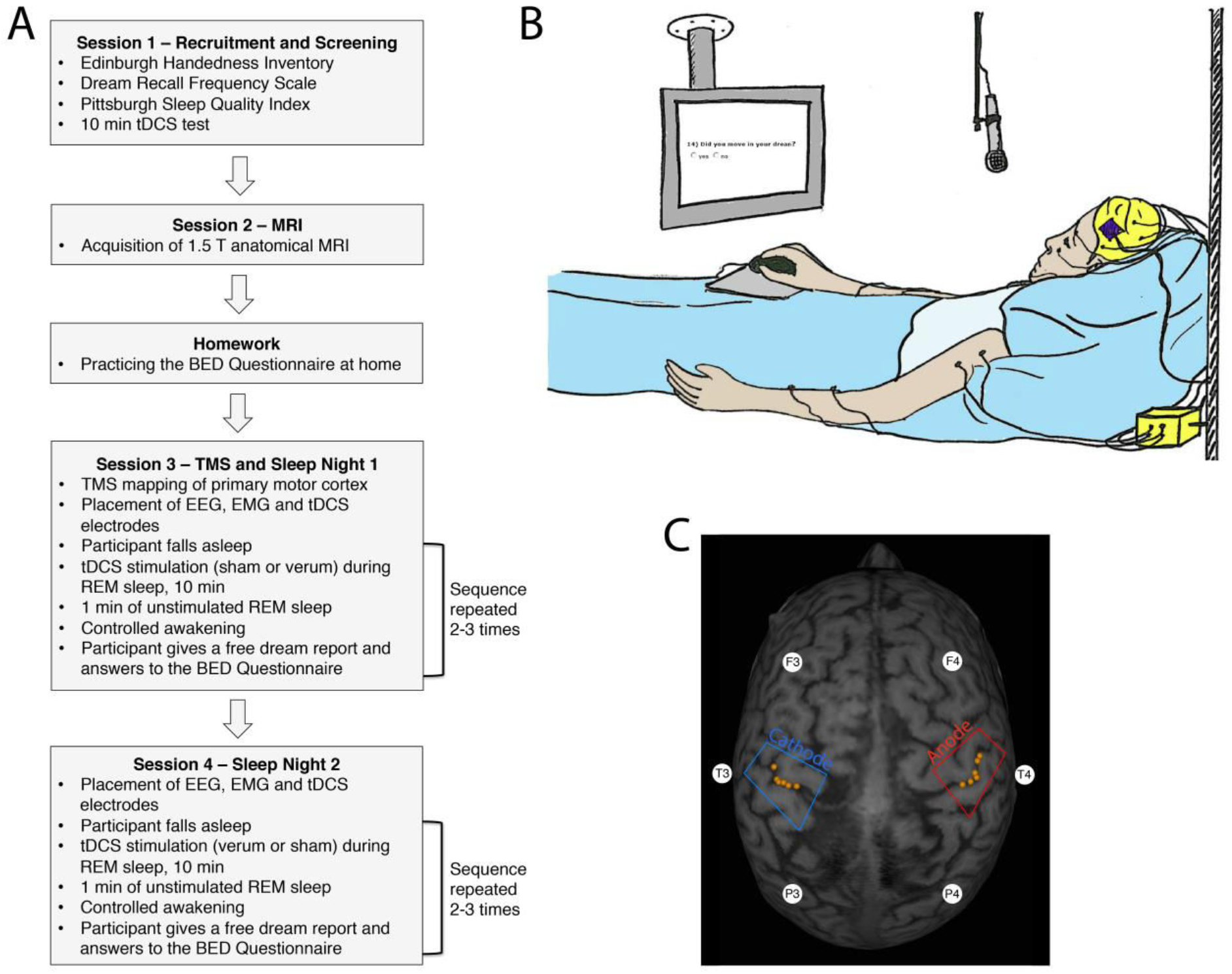
Experimental design. **(A)** Time course of the study. **(B)** Experimental setup during sleep sessions. **(C)** Primary sensorimotor hand areas of a representative participant. Orange dots indicate stimulation sites where TMS pulses induced a subjectively experienced hand movement and/or muscle twitch (located approximately at the central sulcus between the somatosensory and somatomotor cortices). The blue box drawing over the left hemisphere represents the cathode tDCS electrode placement site, and the red box drawing over the right hemisphere represents the anode electrode placement site. White circles depict the approximate location of 6 electrodes used for the EEG inter-hemispheric coherence analysis.

### Participants

Aiming to recruit 10 right-handed individuals with high dream recall frequency and good sleep quality, potential participants were screened with the Edinburgh Handedness Inventory (Oldfield 1971) and the Dream Recall Frequency (DRF) scale (Schredl 2002), which assesses the frequency with which people are able to remember dreams at home. The DRF scale consists of a single question “How often do you remember your dreams?” and 7 possible answers: 0=never, 1=less than once a month, 2=about once a month, 3=twice or three times a month, 4=about once a week, 5=several times a week, and 6=almost every morning. Furthermore, potential participants filled in the Pittsburgh Sleep Quality Index (PSQI) (Buysse et al. 1989). We aimed to recruit individuals whose global PSQI score did not exceed 4 (with 0 indicating no sleep difficulty and 21 indicating severe difficulties in sleep) and whose sleep latency score indicated they typically needed less than 30 minutes to fall asleep.

Given that the application of tDCS may occasionally induce itching, tickling, heat sensations under the electrodes, or even a temporary headache (Priori 2003), we introduced potential participants to the tDCS technique before they made their final commitment to take part in the study. After screening for MRI and tDCS contraindications, they were given the opportunity to familiarize themselves with the tDCS procedure before spending their first night in the laboratory. Participants were stimulated for 10 min with tDCS of 1 mA current over the C3 and C4 electrode sites according to the 10-20 EEG system (approximately over the sensorimotor cortex), which helped them decide whether they wanted to participate in the actual experiment. This also helped minimize the risk that tDCS during REM sleep would lead to awakening.

After screening 16 potential participants, we were able to recruit 10 healthy right-handed university students (4 men and 6 women, mean age 26.8, range 4.4 years). The mean handedness index was 0.9 (SD=0.11; range 0.73 to 1). The mean DRF score was 5.4 (SD=0.79, Min=4, Max=6), indicating high spontaneous dream recall. While this might introduce bias towards high recallers’ dreams, it is arguably the most feasible recruitment strategy for a costly and time-consuming sleep laboratory study. All participants gave their written informed consent according to the Declaration of Helsinki, and the protocol of the study was approved by the Ethics Committee of the Hospital District of Southwest Finland. Participants were financially compensated with 40 euros per night and 10 euros per hour for daytime testing.

### MRI-TMS mapping of the primary sensorimotor hand area

ECoG measurement of the electric field induced by tDCS in a human patient as well as computational modelling of tDCS effects in healthy participants suggest that the spatial focality of tDCS decreases if stimulation electrodes are misplaced by >1cm (Opitz et al. 2018). Thus, aiming to constrain between-participant variance of the stimulation focus below 1cm, the location of the hand area in the primary sensorimotor cortex in both hemispheres was determined individually for each participant with the help of magnetic resonance imaging (MRI) and transcranial magnetic stimulation (TMS). Anatomical brain images were acquired with a 1.5 T MRI scanner Philips Intera at the Turku PET Centre. 3D models of the brain were created using 3D T1-weighted MR sequence. A hospital radiologist confirmed that the brain MRI was normal in all cases. Afterwards, the approximate location of primary sensorimotor hand representations was visually determined from anatomical brain images based on macro-anatomical landmarks (Yousry et al. 1997).

Based on this analysis, the location of the primary sensorimotor hand area was determined for each participant in a separate TMS session, which was carried out on the evening of the first experimental night at the Department of Psychology at the University of Turku. TMS pulses were delivered using eXimia™ TMS stimulator with NBS navigation system (Nexstim Ltd., Helsinki, Finland), which allowed us to navigate within individual anatomical MRI with an approximately 6-mm spatial resolution containing all sources of errors (Ruohonen and Karhu 2010). Participants sat on a reclining chair with their eyes closed and both arms supported by a pillow to ensure that their arm muscles were relaxed. TMS was carried out in a single pulse mode using a figure-of-eight-shaped coil that was held tangentially against the participant’s head. The current direction of the second phase of the biphasic pulse was oriented perpendicularly to the post-central gyrus in the posterior to the anterior direction at the bank between pre-central and post-central sulci (Richter et al. 2013) (see Figure 1B).

First, a rough location of the hand area was estimated by asking participants to report whether they experienced any hand movement following a TMS pulse over the motor cortex in the contralateral hemisphere. Once a reliable hotspot was found, an individual motor threshold, i.e. the minimum TMS intensity required to induce the subjective experience of a hand movement, was determined with the maximum likelihood threshold hunting (MLTH) procedure (Awiszus, 2003). In this process, 20 pulses were delivered to the hand area with different stimulus intensities, starting at 60% of maximal TMS intensity. The mean motor threshold was 56.1% (SD=12.4, Min=28, Max=76.7) of maximal TMS intensity for the left hemisphere, and 59.2% (SD=16.5, Min=24.8, Max=77.12) for the right hemisphere. While motor thresholds did not differ systematically between the hemispheres (paired samples t test: t(9)=1.02, p=0.34, Bf in favor of the null=2.2), there was a strong inter-hemispheric correlation of motor thresholds (Pearson correlation: r=0.82, p=0.004).

Following estimation of individual motor thresholds, the most ventral and caudal points of the hand representation in the primary motor cortex were estimated by delivering TMS pulses with the intensity of 10% above the level of the individual motor threshold. This procedure was consecutively performed for both hemispheres, yielding bilateral hand representation maps that were later used to place tDCS electrodes (see Figure 2B).

**Figure 2.**
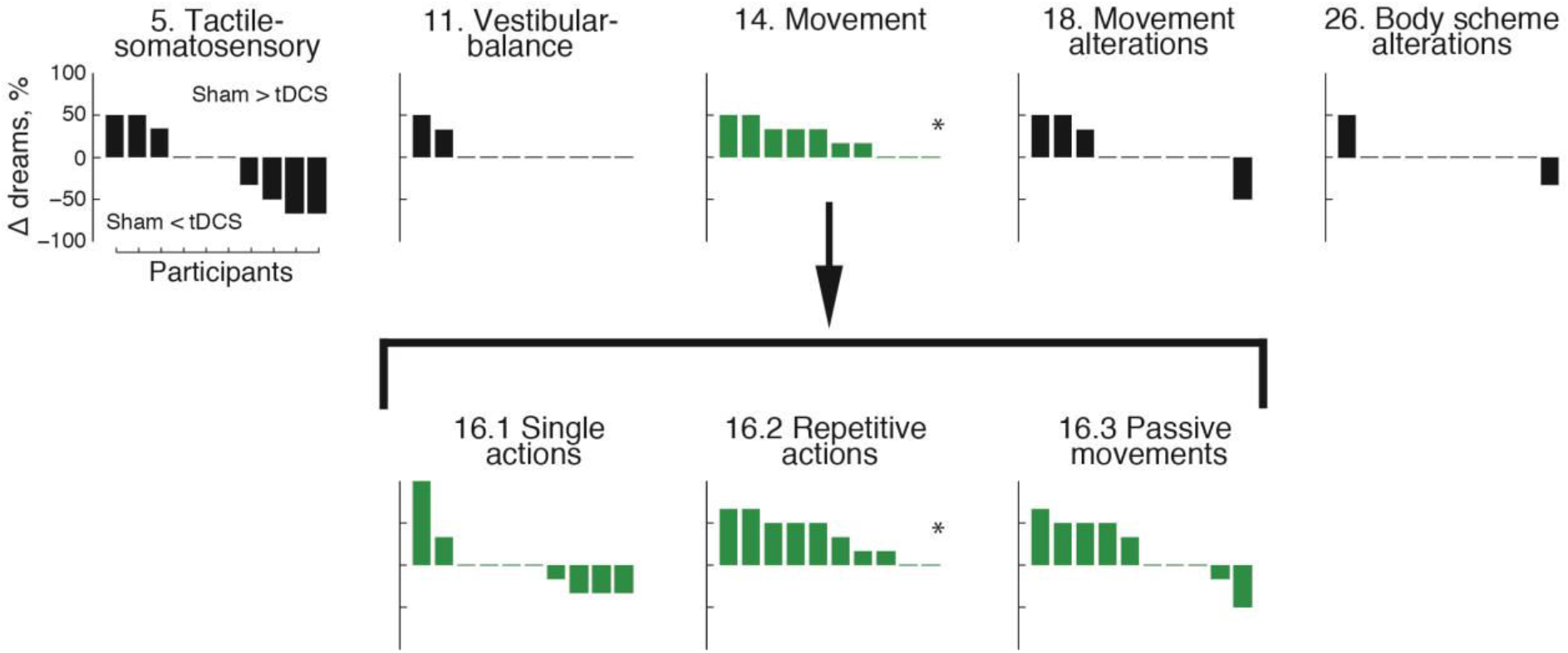
tDCS effects on reported dream experiences. Changes between sham-stimulation and tDCS conditions across the five general categories of dream content (A-E) and for particular kinds of movement (F-H) per participant. Positive and negative values indicate a higher proportion of dreams with a specific experience in the sham-stimulation and tDCS condition, respectively. Individual participants are sorted in descending order beginning with the participant with the highest proportion of dreams with a specific experience in the sham-stimulation condition, compared to the tDCS condition. Participants are sorted separately for each dimension of experience. * p_B_ < 0.05.

### tDCS over the primary sensorimotor cortex during REM sleep

tDCS and sham-stimulation sessions were conducted in the Sleep Laboratory of the Centre for Cognitive Neuroscience at the University of Turku over two non-consecutive nights with each participant. Microprocessor-controlled programmable 1-channel Eldith DC-Stimulator PLUS (Electro-Diagnostic & Therapeutic Systems GmbH, Ilmenau, Germany) was used as a stimulation device.

tDCS was applied bilaterally to the hand area in order to modulate the excitability level of the primary sensorimotor cortex during REM sleep. Participants were asked to avoid caffeine for 6 hours and alcohol and other CNS-affecting drugs for 24 hours prior to the experiment. To ensure these requirements were met, participants filled out the custom-made Pre-Sleep Questionnaire before each session. For each participant, the two stimulation sessions were separated by at least one week in order to avoid interference effects.

Two 35 cm^2^ sized sponge-covered rubber electrodes were soaked with water, and Ten20 electrode paste (Weaver and Company) was applied on both sides of the sponge. The electrodes were placed bilaterally along the central sulcus posterior to the primary motor hand areas, which was determined with the help of MRI-guided TMS (see Figure 1B). They were supported with a comfortable bandage throughout the night. tDCS was carried out on one experimental night and sham-stimulation took place on another night. Participants were blind to the experimental conditions, i.e. whether the tDCS session was followed by the sham session (N=5) or vice versa. An equal number of participants was assigned randomly to each condition.

During the tDCS night, 1mA electric current was delivered to participants’ scalp two or three times per night for 10 min during REM sleep, starting with the second sleep cycle. It has been reported that changes in current direction may result in qualitatively different motor effects, with cathodal stimulation being more effective and largely inhibitory and anodal stimulation being less effective and largely facilitatory (Nitsche et al. 2008). Furthermore, tDCS induced neuroplasticity may accumulate over time (Nitsche and Paulus 2000). In order to keep the stimulation effects consistent throughout the night, the electrode over the right sensorimotor area was always the anode, and the electrode over the left sensorimotor area was always the cathode. This procedure ensured that the asymmetric stimulation during one sleep cycle would not interfere with or cancel stimulation effects during another cycle. We chose to place the cathode over the dominant left hemisphere with the aim to disrupt dream movements.

During the sham-stimulation night, stimulation was conducted by switching on the DC device and stimulating only for 10 sec each at the beginning and end of a 10 min period during REM sleep. Stimulation that lasts only a few seconds has been shown to produce a minimal effect on the brain, if any (Hummel et al. 2005). The aim of sham stimulation was to mimic the skin sensation that is occasionally experienced during the onset and offset of tDCS. This procedure is thought to make the two conditions subjectively indistinguishable (Gandiga et al. 2006). The same procedure was repeated two or three times starting with the second sleep cycle.

### Electrophysiological recordings

To record EEG activity, 16 electrodes (Fp1, Fp2, F7, F3, Fz, F4, F8, T3, T4, T5, P3, Pz, P4, T6, O1, O2) were placed on the scalp following the standard 10-20 system (Jasper 1958). C3, Cz and C4 electrode locations were left empty for the placement of tDCS electrodes. To record eye blinks and vertical saccades, two electrooculography (EOG) electrodes were placed below and above the left eye, while two other electrodes placed adjacent to the lateral canthi of each eye were used to measure horizontal saccades. An electromyography (EMG) electrode placed on the chin was used to record muscle tone, which was used for the scoring of sleep stages. The reference for all these electrodes was placed on the right ear mastoid and the ground electrode was placed on the temple. In addition, two bipolar EMG channels were used to record muscle activity in the right and the left arm flexor digitorum profundus, which were later used to analyze peripheral motor activity. Another two EMG channels recorded activity of the deltoid muscles in both arms. Electrophysiological recordings were continuously monitored on a computer screen and all electrodes were regularly checked throughout the night to ensure that the impedance remained under 5 kΩ. All data were recorded at 500 Hz sampling rate with Ag/AgCl electrodes using NeuroScan amplifier SynAmps Model 5083. Given that tDCS onset induces a slow frequency artifact in the EEG that may preclude online polysomnographic scoring, a 1-Hz high-pass filter was applied during recording for online monitoring of sleep stages (Marshall 2004). As expected, tDCS onset- and offset-induced artifacts always faded away after 5-10 sec.

### Collection of dream reports

One minute after the termination of tDCS or sham-stimulation, participants were awakened from REM sleep with a standard awakening tone. They were then asked to give a verbal dream report of “everything that was going through their minds before awakening”, aiming to facilitate dream recall. Afterwards, participants were asked if they remembered anything else about their dream. To avoid a possible bias between stimulation conditions, these questions were played from a pre-recorded computer audio file. Following the free dream report, participants were asked to fill in the Bodily Experiences in Dreams (BED) Questionnaire. The questionnaire was designed as an internet survey programmed on www.webropol.com and was projected on a screen above the bed in the sleep laboratory. Participants navigated and responded to the BED Questionnaire by controlling a mouse while lying in bed.

Participants were stimulated and awakened two or three times per night, depending on how many REM sleep periods they had. The number of awakenings was balanced across the first and the second night and across the two stimulation conditions (see Table S3). White dream reports (i.e. cases when a person reports the occurrence of dream experiences but cannot recall any specific details) as well as sleep mentation reports (i.e. when a person reports non-perceptual subjective experiences, such as thinking) were excluded from the analysis. A total of 50 dreams reported during a total of 20 nights were available for analyses.

### Bodily Experiences in Dreams (BED) Questionnaire

The 41-item BED Questionnaire was designed to gather detailed information about kinaesthetic and other bodily experiences in dreams (see Appendix 1). The BED Questionnaire consists of 5 general questions with respective sub-scales (see Table 1). Each of the general questions targets a particular category of body-related experience: vestibular sensations, tactile and somatosensory experiences, movement, movement alterations, and body schema alterations. Each general question, if answered positively, is followed by sub-scales targeting more specific instances of this category of experience. For example, if a participant indicated that they had experienced movement sensations, they would then be asked about the occurrence of specific types of movement sensations, such as single, repetitive, and passive movements. In addition, participants were asked whether the reported sensation concerned the whole body, the right or left hand, the right or left side of the face, or another body part (see Appendix 1). If they answered negatively, they would skip to the next general category. Depending on whether a sub-scale asked about the intensity or the duration of experience, 9 point Likert-scales for answering ranged either from “1=Low intensity” to “9=High intensity” or from “1=Never” to “9=Throughout”.

**Table 1.**
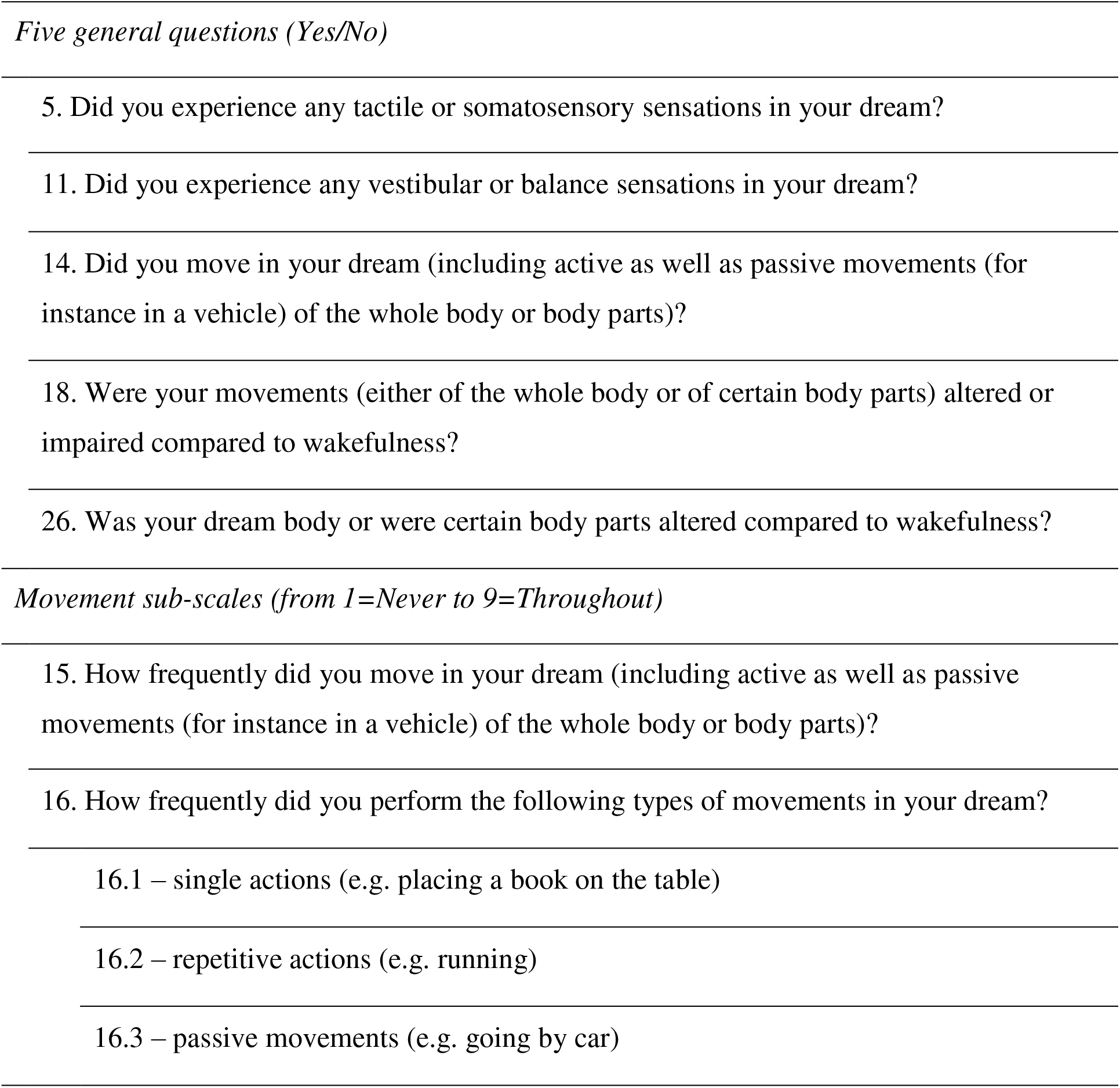
The BED Questionnaire: General questions and exemplary sub-scales

### Word count of verbal dream reports

The length of dream reports was assessed by two blind judges (authors JW and KV), who independently calculated the meaningful word count of each dream report. Murmurs, repetitions of words, and any secondary reflections or comments about the dream were not included in the word count. The judges initially agreed on the word count of 47 out of 50 dream reports (94% agreement). The judges discussed the reasons for the mismatch in the remaining 3 cases and reached an agreement.

### Content analysis of movement sensations in verbal dream reports

Following the findings from the BED Questionnaire, we carried out content analysis of verbal dream reports, focusing on the specific types of movement (single actions, repetitive actions, passive movement) performed by the dream self. To compare the type and frequency of movements reported in the BED Questionnaire to those explicitly mentioned in dream reports, two blind judges (authors VN and BL) carried out a content analysis of verbal reports. First, the judges scored whether each dream report contained at least one movement produced by the dream self, excluding facial movements such as talking, drinking, and blinking, as we reasoned that individuals do not typically consider facial musculature when asked to report their movements. Movements attributed to the first-person plural “we” were treated as involving movements of the dream self. Second, the judges identified individual movements produced by the dream self in each dream that, in the first step, was judged to contain movement. Third, they scored the type of the identified movements (single action, repetitive action, passive movement). All three stages of the content analysis were first carried out individually and the obtained results were then compared between the judges. In the case of disagreement, the judges discussed it until an agreement was achieved.

Regarding the presence or absence of movement in a given report, the judges initially agreed on 45 out of 50 dream reports (90% agreement). After discussion, the judges agreed that the remaining 4 dreams contained references to movements produced by the dream self, while one report had no explicit references to such movement. Regarding individual movements, judges initially agreed on the identification of 33 movements, and disagreed on 19 movements (63.5% agreement). The disagreement was caused by one judge either missing a movement or treating it as part of a longer sequence of movements, e.g. treating walking from A to B and from B to C as a single movement. After discussion, the judges agreed that the dream reports contained a total of 48 individual movements executed by the dream self. Regarding specific types of movements (single action, repetitive action, passive movement), the judges initially agreed on 44 out of 48 movements (91.7% agreement). After discussion, the judges agreed that the remaining 4 movements should be scored as follows: “diving” - single action, “riding a bike downhill” - passive movement, “writing something” - repetitive action, “made some coffee” - single action.

### EEG analysis: coherence and spectral power

To assess the electrophysiological effects of tDCS on brain functioning, we analyzed the full period of 1 min of EEG signal recorded between the termination of tDCS or sham-stimulation and controlled awakening. tDCS artifacts did not contaminate this EEG interval whilst sleep scoring ensured that REM sleep continued up to the point of awakening. On one occasion, a spontaneous awakening took place before the planned controlled awakening, and only 7 sec of stimulation-free sleep EEG were available for analysis. On another occasion, a spontaneous awakening took place immediately after the termination of stimulation; this recording was excluded, leaving 49 EEG recordings available for analysis.

Continuous recordings were first high-pass (0.5 Hz) and then low-pass (45 Hz) filtered using a FIR filter as implemented in EEGlab toolbox (Delorme and Makeig 2004). The data were then common average referenced, and excessively noisy periods of recording were manually deleted (an average of 743 ms per single recording). Detached or excessively noisy channels were deselected (an average of 0.2 channels per dataset), and an independent component analysis (ICA) was carried out on the remaining channels, using EEGlab toolbox (Delorme and Makeig 2004). Independent components reflecting eye movements and other sources of noise were manually deleted (an average of 3.3 ICs per recording), following which dropped noisy EEG channels were interpolated using spherical spline interpolation. Continuous EEG recordings were epoched into 2-sec segments with a 50% overlap between adjacent segments. Several epochs that still contained visible artifacts were manually deleted (an average of 0.5 epochs per recording). Individual epochs were demeaned across the whole 2 sec interval.

We analyzed EEG inter-hemispheric coherence in the beta oscillation range (15-30 Hz) at the electrodes adjacent to the stimulation site (F3, F4, T3, T4, P3, P4). Magnitude-squared coherence was computed in the range from 2 Hz to 44 Hz with a maximum frequency resolution of 2 Hz between pairs of EEG channels adjacent to the stimulation site from the frontal (F3-F4), temporal (T3-T4) and parietal (P3-P4) side, using Brainstorm toolbox (Tadel et al. 2011). Coherence values obtained at a single 2 sec segment level were averaged across beta frequency range (15.6-29.3 Hz). Next, coherence values were averaged across each 1-min pre-awakening recordings. Afterwards, individual means were averaged over several awakenings for each participant according to the experimental condition, yielding 10 tDCS and 10 sham-stimulation values for each electrode pair.

In the case of a significant difference between tDCS and sham-stimulation conditions across the 1-min pre-awakening periods, coherence was computed at four separate 15 sec sub-intervals preceding controlled awakening: −60 to −46 sec, −45 sec to −31 sec, −30 to −16 sec, and −15 to −1 sec. A significant difference between tDCS and sham conditions observed immediately after the termination of stimulation (−60 to −46 sec) was expected to reflect a tDCS-driven modulation of EEG activity, as an effect size of neurophysiological changes following motor tDCS decreases with increasing time (Nitsche and Paulus 2000). Contrary to this, a significant difference between tDCS and sham-stimulation conditions at the interval preceding awakening (−15 to −1 sec) with no difference at the −60 to −46 sec interval was expected to reflect an unspecific modulation of EEG activity, e.g. micro-arousals caused by tingling sensations could eventually trigger body movements in bed.

To control for a possible confound of EEG spectral power on coherence computation (Bowyer, 2016), we carried out a control analysis of EEG beta power. Spectral power was computed across 2 sec epochs using Hilbert transform, set from 1 Hz to 44 Hz in steps of 1 Hz, for the same set of 6 electrodes adjacent to the stimulation site. Power values obtained at a single 2 sec segment level were averaged across beta frequency range (15-30 Hz), with subsequent data averaging steps repeating coherence analysis.

### Phasic EMG analysis

We investigated the effects of tDCS on peripheral muscle tone by analyzing EMG activity from the left/right arm flexor and deltoideus muscles during the 1 min interval between the termination of tDCS or sham-stimulation and the controlled awakening of participants. We were specifically interested whether EMG traces following tDCS and sham-stimulation showed increased phasic muscle activity compared to the pre-stimulation baseline window, and whether bihemispheric tDCS modulated interaction between the left/right arm EMG. Since phasic EMG activity manifests during REM sleep as short-lasting muscle bursts recorded by surface electrodes (Fairley et al. 2012), we split the 1-min epochs into 60 non-overlapping 1-sec segments and carried out a binary assessment whether each segment contained phasic EMG activity. Segments with phasic EMG activity were then assigned a value of one, segments without phasic EMG activity a value of zero. The mean overall 60 binary values were then used to define the ratio of phasic EMG activity within the respective epoch.

More specifically, since phasic EMG activity is reflected in broadband spectral power changes, we used the variance of gamma band (50-250 Hz) power for the detection of short-lasting bursts of muscle activity. In a first step we high-pass filtered the raw data with a 3rd order butterworth filter with a cutoff-frequency at 50Hz (Suppl. Fig. 1 a-b). For the subsequent time-frequency analysis, we used a single-tapered spectral analysis method (Percival and Walden 2000) with a time window of 50 ms and 10-ms time steps. The relative power changes were then calculated by dividing the time-resolved amplitude for each frequency bin by the frequency-specific average of the whole 1 min epoch (Suppl. Fig. 1 c). After splitting the epochs in 1-s segments, the variance of relative power was calculated for each segment and every frequency. The variance of gamma band power was then defined as the mean over all frequencies between 50 Hz and 250 Hz.

To assess a relative shift towards more phasic/tonic activity in response to stimulation, the variance of gamma band power was calculated both for the 60 sec epochs after the termination of tDCS or sham-stimulation and for a 30 sec baseline time window before tDCS or sham-stimulation. The relative variance of gamma band power was then calculated by dividing the variance by the averaged variance in the baseline time window (Suppl. Fig. 1 d). This way, post-stimulation segments with the variance of gamma band power higher than the corresponding average (median) in the stimulation-free 30 sec baseline time window received a relative variance value greater than one and were defined as segments shifting towards phasic EMG, while segments with a relative variance between zero and one were defined as segments shifting towards tonic EMG. Finally, a proportion of phasic segments was calculated across the whole 60 sec post-stimulation epoch, yielding values ranging from 0, indicating a complete shift towards tonic EMG, to 1, indicating a complete shift towards phasic EMG.

### Statistical analysis

All dependent measures were averaged per individual participant separately for the sham-stimulation and tDCS conditions. A Shapiro-Wilk test was used to assess the distribution normality of dependent variables. Paired-samples t test and Pearson correlation were carried out when distribution of given variables (or their difference in a case of t test) was normal, and Wilcoxon signed-rank test (Z statistic) and Spearman rank order correlation were used in the cases of non-normal distribution of one or both variables. For the paired-samples t-test, Cohen’s *d* was calculated as an effect size estimate using pooled variance. For the Wilcoxon signed-rank test, r=Z/sqrt(N) was calculated as an effect size estimate. All statistical tests were two-tailed. To control for multiple comparisons, Bonferroni correction was applied by multiplying the obtained p value by the number of comparisons with a given set of tests. Bonferroni corrected p values are denoted as p_B-N_ where N indicates the number of multiple comparisons. For all control analyses, we report uncorrected p values. For the control t tests where we expected null findings, we additionally report Bayes factor in favor of the null. Statistical analyses were carried out with SPSS 22 and JASP 0.8.2.

## Results

### tDCS modulates dream movement

The first research question addressed whether the sensorimotor cortex is involved in the generation of bodily experiences in dreams. To answer this, we compared the percentage of dreams with different types of bodily experiences reported in the BED Questionnaire between the tDCS and sham stimulation. Among the general dimensions of bodily experience in dreams (tactile/somatosensory, vestibular/balance, movement, movement alterations, body scheme alterations), we found a significant difference only for movement (see Fig 2 and Table 2). Specifically, the proportion of dreams with movement was significantly lower in the tDCS (M=63.1%, SEM=10.2%) compared to the sham-stimulation (M=86.6%, SEM=7%) condition (paired samples t test: t(9)=3.77, p_B-5_=0.022, d=0.85). That is, participants were less likely to answer YES to the question “Did you move in your dream?” when they were awakened 1 min after termination of bihemispheric tDCS. At the individual level, 7 out of 10 participants showed this effect, whereas the remaining 3 participants had equal proportions of dreams with movements between the two conditions (see Fig. 2).

**Table 2.** The BED Questionnaire: Percentage of dream reports containing specific bodily experiences following sham-stimulation and tDCS during REM sleep

| *Bodily experiences* | *Sham* | *tDCS* | *Statistical test* |  |
| --- | --- | --- | --- | --- |
|  | *M (SEM)* | *M (SEM)* | *t/Z* | *p<sub>B</sub>* |
| *Five general dimensions* |  |  |  |  |
| 5. Tactile- somatosensory | 34.9 (12.3) | 43.2 (12) | t(9) = 0.59 | 1 |
| 11. Vestibular- balance | 8.3 (5.7) | 0 (0) | Z = 1.34 | 0.9 |
| 14. Movement | 86.6 (7) | 63.1 (10.2) | t(9) = 3.77 | 0.022* |
| 18. Movement alterations | 13.3 (6.9) | 5 (5) | Z = 0.76 | 1 |
| 26. Body scheme alterations | 5 (5) | 3.3 (3.3) | Z = 0.45 | 1 |
| *Movement sub-scales* |  |  |  |  |
| 16.1 Single actions | 53.3 (13.3) | 51.7 (13.3) | Z = 0.22 | 1 |
| 16.2 Repetitive actions | 65 (9.8) | 30 (8.5) | t(9) = 4.36 | 0.006* |
| 16.3 Passive movements | 30 (8.5) | 11.7 (7.9) | t(9) = 1.56 | 0.45 |

*Note.* t: paired samples t test; Z: Wilcoxon signed-rank test; \* p<sub>B</sub>< 0.05

To investigate whether specific types of movement were inhibited by tDCS, we compared the proportion of dreams with single actions (i.e. movements that are not repeated immediately after their execution, such as placing a book on the table), repetitive actions (i.e. the same movements repeated several times in a continuous sequence, such as running), and passive movements (i.e. movements determined by external forces, such as traveling by car) between tDCS and sham-stimulation conditions (see Table S1 for examples of movement descriptions in the verbal dream reports). There were significantly less dreams with repetitive actions in the tDCS condition (M=30%, SEM=8.5%) compared to the sham condition (M=65%, SEM=9.8%) (paired samples t test: t(9)=4.36, d=1.21, p_B-3_=0.006) (see Fig. 2). There were no significant tDCS effects on the frequency of dreams containing single actions or passive movements (see Table 2).

Interestingly, we found no difference in movement frequency between the stimulation conditions in verbal dream reports that were content analysed by external judges (see Table S2). This could be due to a considerably smaller proportion of explicitly expressed movements in free reports compared to the BED Questionnaire answers. It is possible that participants tended to omit movements from the spontaneous verbal reports that were given before answering to explicit motor questions of the BED Questionnaire (see Supplementary Results).

According to our questionnaire data, a majority of dream movements involved the whole body (M=75.5%, SEM=7.62%) and more rarely the right hand (M=25%, SEM=8.23%) or both hands (M=15.83%, SEM=7.02%); another unspecified body part was mentioned in only one report. Repetitive actions typically involved the whole body (M=89.8%, SEM=6.8%), with only 5.6% of repetitive movements performed by the right hand (Wilcoxon signed-rank test: Z=2.71, p=0.007, effect size *r*=0.64). Contrary to this, the proportion of single actions was comparable for the whole body (M=43.8%, SEM=12.3%) and right hand movements (M=34.4%, SEM=11.5%, Wilcoxon signed-rank test: Z=0.43, p=0.67, effect size *r*=0.11). No systematic body part or laterality differences were observed between the sham-stimulation and tDCS conditions.

Importantly, the observed reduction of dream movement following tDCS was not related to the overall length of dream reports, which could have been a confounding factor. To test whether the reduction in dream movement was related to shorter dream reports following tDCS, we compared the subjectively reported duration of dreams during the tDCS and sham-stimulation conditions (BED Questionnaire - Q41, see Appendix 1). There was no difference in the subjectively reported duration of dream reports between tDCS (Median=9.17 min, range from 1.5 min to 97.5 min) and sham-stimulation (Median=9.67 min, range from 0.83 min to 40 min) conditions (Wilcoxon signed-rank test: Z=0.36, p=0.72, r=0.11). Furthermore, we compared the word count of dream reports. Once again, there was no significant difference between tDCS (M=76.1, SEM=16.31) and sham-stimulation (M=124.2, SEM=34.68) conditions (paired samples t test: t(9)=1.69, p=0.124, d=0.56, Bf in favor of the null=1.11). On four occasions, participants remembered and reported additional details of a dream after completing the original dream report and questionnaire, while they were trying to fall asleep again. When these secondary reports were included in the word count analysis, there was still no significant difference in word count between tDCS (M=89, SEM=19.83) and sham-stimulation (M=124.98, SEM=34.55) conditions (paired samples t test: t(9)=1.15, p=0.281, d=0.4, Bf in favor of the null=1.91). We thus conclude that differences in the length of dream reports (and in the subjectively estimated duration of dreams) were not related to the observed reduction of dream movement following tDCS.

### tDCS modulation of EEG activity

Given the opposing direction of bihemispheric tDCS in the current study, i.e. a cathodal inhibitory effect over the left motor cortex and anodal excitatory effect over the right motor cortex, we hypothesized that a reduction of repetitive whole-body actions in response to tDCS was due to a decreased inter-hemispheric coordination of motor processing. To investigate this hypothesis, we restricted EEG analysis to the beta frequency band, because (1) transient and tonic changes in EEG beta oscillatory activity underlie cortical processing of both real (Gerloff et al. 1998; Jenkinson and Brown 2011; Zaepffel et al. 2013) and imagined (Neuper et al. 2005; Nam et al. 2011) movements, (2) preparation and execution of movement involves inter-hemispheric functional coupling in the beta frequency range (Leocani et al. 1997; Mima et al. 2000), and (3) motor impairment and successful rehabilitation involve changes in the inter-hemispheric interaction in the beta frequency range (Pellegrino et al. 2012; Fortuna et al. 2013). We thus expected bihemispheric tDCS to destabilize motor processing by reducing inter-hemispheric coherence in the beta frequency range.

As predicted, we observed a significant decrease in coherence between parietal electrodes P3-P4 following tDCS compared to sham-stimulation during a 1-minute stimulation-free period before awakening (Wilcoxon signed-rank test: Z=2.5, p_B-3_=0.039, effect size *r*=0.79). No inter-hemispheric tDCS effects were observed between frontal (paired samples t test: t(9)=0.72, p_B-3_=1, d=0.244) or temporal electrodes (t(9)=0.38, p_B-3_=1, d=0.114). To control for temporal specificity of the decrease of parietal coherence, we repeated the same analysis in four separate time intervals following termination of stimulation: −60 to −46 sec, −45 to −31 sec, −30 to −16 sec, and −15 sec to −1 sec prior to awakening. A significant effect observed only in the time window before awakening (i.e. −30 to −16 sec, and/or −15 sec to −1 sec) would indicate a non-specific effect of experimental stimulation. Compared to sham-stimulation, a significant decrease of parietal coherence took place in the tDCS condition throughout all four sub-intervals between the offset of stimulation and the onset of awakening, confirming a direct and relatively long-lasting tDCS effect on parietal coherence in the beta-frequency range (see Fig 3).

**Figure 3.**
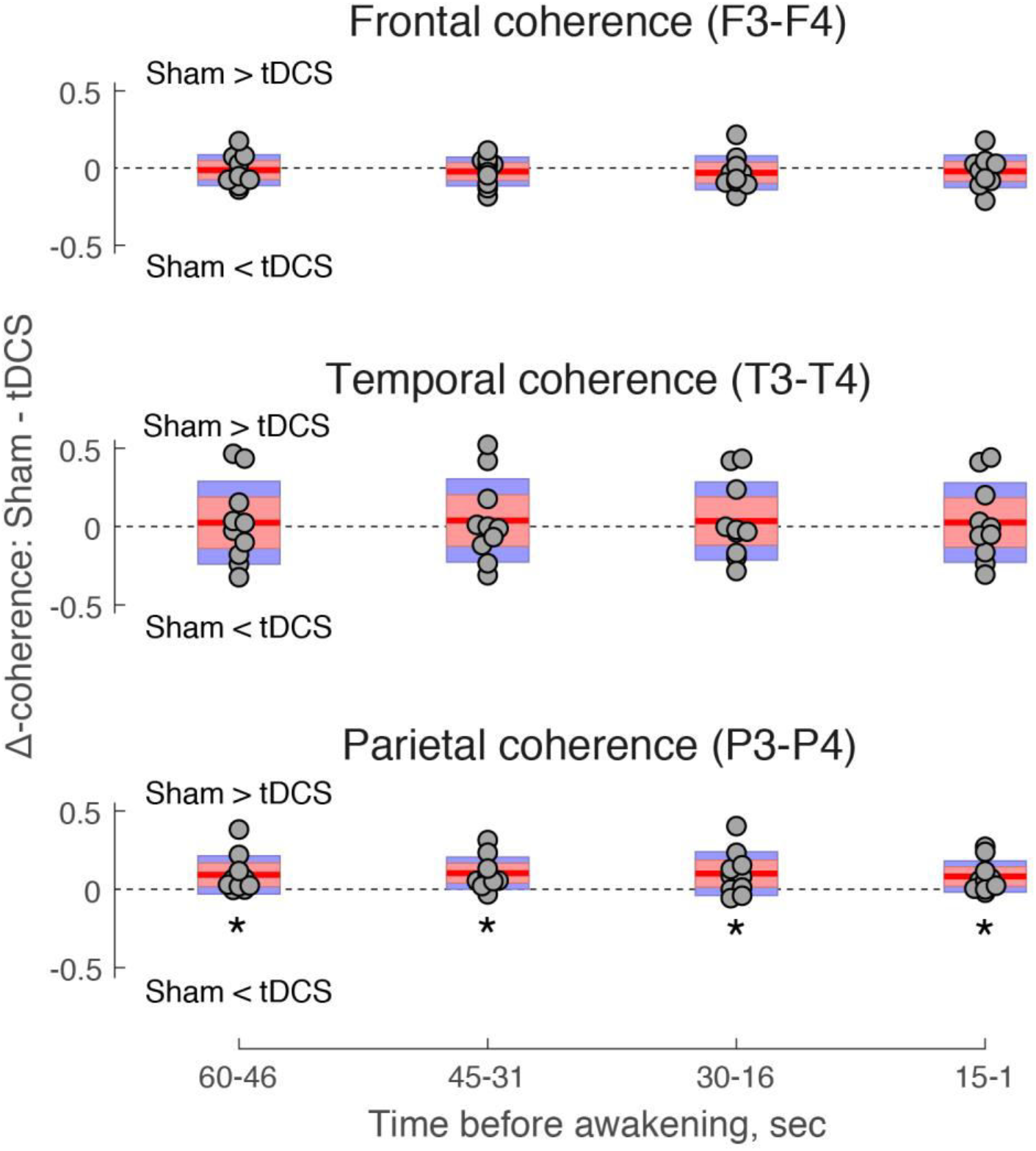
EEG coherence following tDCS during REM sleep. Inter-hemispheric EEG coherence between frontal (top), temporal (middle), and parietal (bottom) electrodes surrounding the tDCS site, expressed as a difference between sham-stimulation and tDCS conditions (Δ-coherence). Jittered circles represent individual participants. Red lines depict the mean of Δ-coherence, pink bars represent 1 standard deviation (SD), and blue bars represent 95% confidence intervals for the mean. Positive values indicate higher coherence in the sham-stimulation condition, whereas negative values indicate higher coherence in the tDCS condition. Δ-coherence is plotted separately in four stimulation-free time intervals preceding controlled awakenings from REM sleep. In the parietal region, coherence was reduced by tDCS compared to sham stimulation in −60- to 46 sec (Z=2.5, p=0.013, r=0.79), − 45 to −31 sec (t(9)=3.17, p=0.011, d=0.97), −30 to −16 sec (t(9)=2.27, p=0.05, d=0.88) and −15 to −1 sec (t(9)=2.57, p=0.03, d=0.74) time intervals. * p < 0.05.

Given that EEG coherence can be affected by spectral power differences between conditions (Fein et al. 1988), we carried out a control analysis to compare beta power in the electrodes adjacent to the stimulation site across a 1 min stimulation-free pre-awakening period. There was a significant decrease of beta power at the left parietal site (P3) in the tDCS compared to the sham-stimulation condition (paired samples t test: t(9)=2.29, p=0.048, d=0.37, Bf in favor of the null=0.64), whereas tDCS did not modulate beta power in the right parietal site (P4) (t(9)=0.73, p=0.48, d=0.088, Bf in favor of the null=2.93). The observed trend was investigated further across four 15 sec sub-intervals. No tDCS effects were observed regarding beta power in P3 electrode during time intervals immediately following motor cortex stimulation, i.e. −60 to −46 sec (paired samples t test: t(9)=1.159, p=0.276, d=0.25, Bf in favor of the null=1.89) and −45 to −31 sec (t(9)=1.172, p=0.271, d=0.3, Bf in favor of the null=1.87). Contrary to this, beta power decreased during time intervals preceding awakenings: −30 to −16 sec (paired samples t test: t(9)=2.433, p=0.038, d=0.433, Bf in favor of the null=0.46) and −15 to −1 sec (t(9)=2.829, p=0.02, d=0.379, Bf in favor of the null=0.28). Given that EEG beta coherence was modulated by tDCS across all four time intervals, we conclude that its decrease was not due to the temporally constricted changes in beta spectral power.

### tDCS modulation of EMG activity

We observed a significant association in the proportion of phasic EMG activity in the flexors between the left and right arms during the 1-min period of REM sleep from the offset of tDCS to the controlled awakening (Pearson correlation: forearm flexors: r=-0.769, p_B-4_=0.037; deltoids: r=-738, p_B-4_=0.06). The negative correlation between the arms likely reflects the asymmetrical modality of stimulation with the cathode placed over the right sensorimotor cortex and the anode over the left sensorimotor cortex. Contrary to this, there was no association in the proportion of phasic EMG between forearms following sham stimulation (Pearson correlation: forearm flexors: r=0.095, p_B-4_=1; deltoids: r=0.308, p_B-4_=1), indicating that muscle activity varied independently (see Fig 4A). Regarding absolute EMG values, there was no difference between phasic activity in the left as compared to the right arm in either the sham-stimulation condition (paired samples t test: forearm flexors: t(9)=0.12, p_B-4_=1, d=0.08; deltoids: t(9)=1.52, p_B-4_=0.66, d=0.57) of following tDCS (forearm flexors: t(9)=1.88, p_B-4_=0.37, d=1.13; deltoids: t(9)=1.96, p_B-4_=0.33, d=1.08).

**Figure 4.**
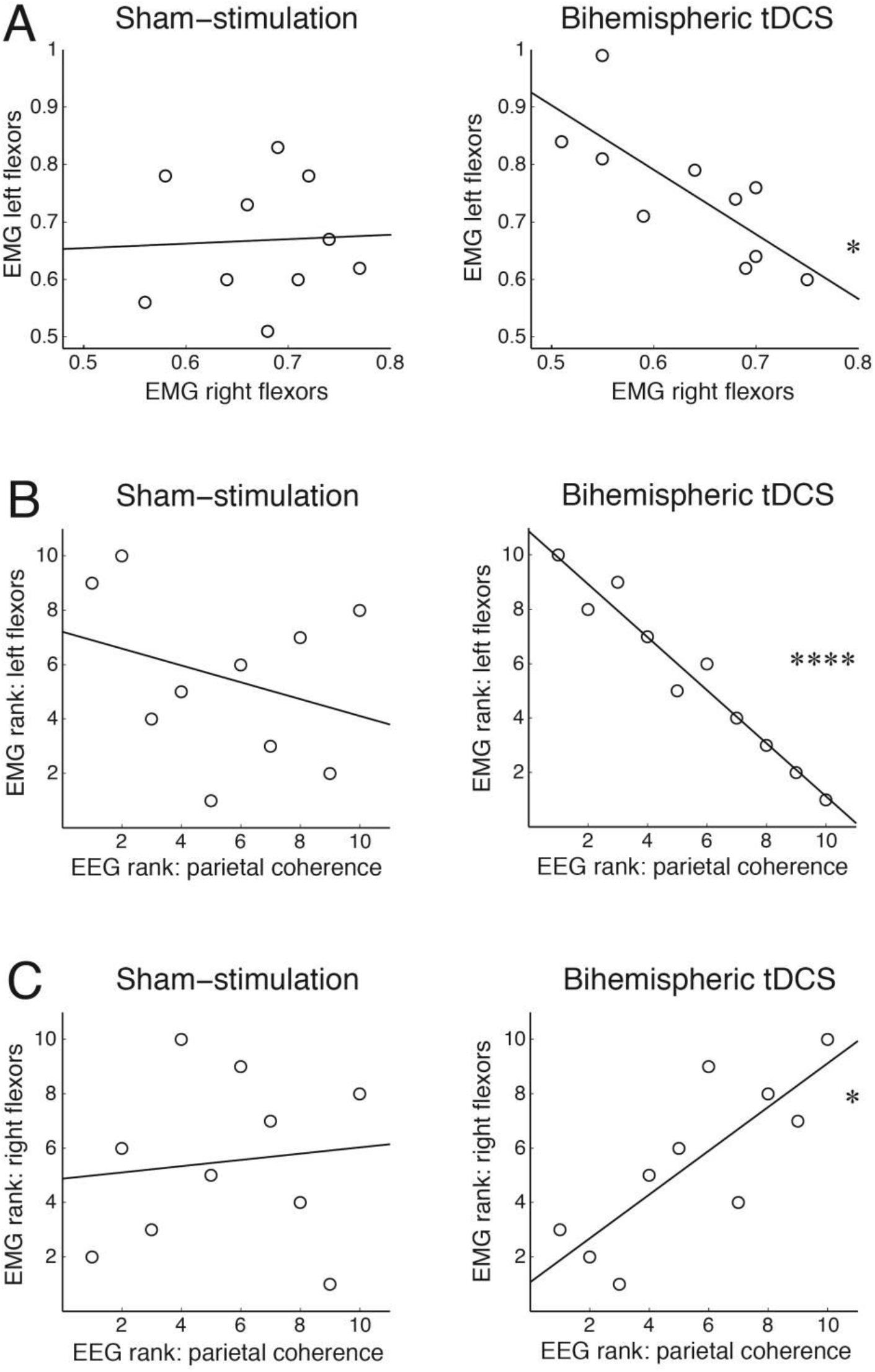
Bihemispheric tDCS during REM sleep modulates phasic activity of the forearm muscles. **(A)** Correlation of EMG shift towards phasic activity between the left and right forearm flexor muscles in the sham-stimulation and tDCS conditions. **(B-C)** Correlation between EMG shift towards phasic activity and EEG parietal coherence in the beta frequency band, plotted separately for the left and right forearm recordings, in the sham-stimulation and tDCS conditions. Ranked data are plotted in (B) and (C) as Spearman’s rank order correlations were carried between EMG and EEG measures. In all plots, the least-squares lines are plotted to visualize associations between variables. * p_B_ < 0.05, **** p_B_ < 0.00005.

Next, we investigated whether peripheral EMG activity is associated with EEG parietal coherence in the beta frequency band, which decreased in response to tDCS during REM sleep (see Fig 4B-C). In the tDCS condition, EEG coherence was significantly associated with the proportion of phasic activity in the left flexor muscles (Spearman rank order correlation: rho=-0.976, p_B-8_=0.00001), and the right flexor muscles (rho=0.806, p_B-8_=0.039). Interestingly, higher parietal coherence was associated with a larger proportion of phasic activity in the right forearm muscles and a lower proportion of phasic activity in the left forearm muscles, once again likely reflecting differential effects of anodal vs. cathodal stimulation. No association was observed between parietal EEG coherence and the proportion of phasic activity in flexor muscles in the sham stimulation condition (lowest p_B-8_=1). Likewise, there was no association between parietal EEG coherence and deltoid EMG, neither during sham-stimulation (lowest p_B-8_=1) nor tDCS conditions (lowest p_B-8_=0.72), indicating a site specific interaction between EEG and EMG measures.

## Discussion

The foremost aim of our study was to investigate the role of the sensorimotor cortex in generating bodily sensations in REM sleep dreams by modulating the excitability of the sensorimotor cortex with tDCS. We found that compared to sham stimulation, bihemispheric tDCS over the sensorimotor cortex reduced the frequency specifically of repetitive actions of the dream self in preceding REM sleep dreams, providing causal evidence that the sensorimotor cortex is involved in the generation of dream movement. Furthermore, tDCS interfered with inter-hemispheric EEG coherence and peripheral EMG activity, pointing to a change in both the central and peripheral motor systems in response to bihemispheric tDCS during REM sleep.

### Frequency of bodily sensations and movement in dreams

To systematically assess and directly interfere with bodily sensations in dreams, we developed a questionnaire designed to capture various dimensions of bodily experiences in dreams (see Table 2 and Appendix 1 for the complete questionnaire). Interestingly, independently of tDCS, our data suggest that while dream movements were very common, other bodily sensations such as somatosensory sensations, vestibular sensations or body schema alterations were rather rare. This overall pattern of frequent dream movement coupled with rare reports of other bodily sensations has been found in previous studies (Hobson 1988; Schwartz 2000; Windt 2018). Our study extends the previous work based on spontaneous dream reports by showing that when different types of bodily experiences are specifically investigated through use of a questionnaire, movements and tactile sensations remain the predominant dimensions of bodily experience in dreams. Thus, content analysis- and questionnaire-based studies provide converging evidence for the important role of sensorimotor phenomena in dreams.

The predominance of dream movement in our data also seems to be in line with a recent suggestion that kinesthesia is central to the generation of dream experience, at least during sleep onset (Nielsen 2017). At the same time, in our study, 36.9 % of dream reports following tDCS contained no movements. It therefore seems that specifically self-movements are not strictly necessary to sustain REM sleep dreaming. Moreover, the decrease of dream movement did not reduce the length of dream reports in our sample. Whether these dreams still involved e.g. observed movement is an open question.

### Electrophysiological effects of bihemispheric tDCS

Bihemispheric tDCS over the sensorimotor cortex, as compared to sham stimulation, specifically altered repetitive actions in dreams. Repetitive actions are typically dependent on implicit memory of learnt motor sequences (e.g., walking), the automatic processing of which does not require explicit awareness and monitoring of movements. Such learnt, automatic movements, as compared to more controlled and deliberate movements, are also associated with a smaller increase of activity in brain areas related to motor processing (Wu and Hallett 2005). Thus, arguably, a relatively modest tDCS interference with cortical processing might have down-regulated motor cortex activity involved in the processing of automatic movements, reducing it to the baseline resting level and simultaneously inhibiting the occurrence of repetitive actions in dreams. Contrary to this, the relatively stronger cortical activation underlying single controlled actions might not have been reduced sufficiently by tDCS interference to significantly alter dream content. This would explain why our results showed a specific decrease in repetitive actions, while the frequency of single actions in dreams remained relatively high during tDCS and did not significantly differ from sham stimulation. Alternatively, bihemispheric stimulation might have interfered with the temporal coordination of dream movement, prohibiting long sequences of repetitive actions, but sparing temporally restricted single actions. Indeed, dream imagery is notoriously unstable and prone to change in discontinuous jumps (Revonsuo and Salmivalli 1995). Such possibilities should be more directly assessed in future studies, e.g. using motor imagery tasks during wakefulness that would allow for a more stringent control of movement complexity.

We found that bihemispheric tDCS interfered with neural processing in the beta frequency band, classically linked to motor processing (Leocani et al. 1997; Gerloff et al. 1998; Mima et al. 2000; Neuper et al. 2005; Jenkinson and Brown 2011; Nam et al. 2011; Pellegrino et al. 2012; Fortuna et al. 2013; Zaepffel et al. 2013; Khanna and Carmena 2015). In our setup, bihemispheric tDCS reduced inter-hemispheric coherence of parietal beta oscillations. Arguably, the differential montage of tDCS electrodes, i.e. the excitatory anode over the right sensorimotor cortex and the inhibitory cathode over the left sensorimotor cortex, disrupted inter-hemispheric coordination of motor commands, reducing the rate of repetitive actions associated with whole body movements in dreams. A differential effect of bihemispheric tDCS was also observed in the phasic EMG activity of the arm muscles. While phasic EMG varied independently between the arms during sham stimulation, a strong negative correlation was observed following tDCS, i.e. it suppressed phasic muscle activity in one arm while increasing it in the other arm.

We expected that such destabilizing and hemisphere-specific effects of tDCS would also cause unilateral distortions of bodily sensations in dreams, i.e. stronger effects on one side of the dream body. However, the observed reduction of dream movement in dreams was independent of the laterality of stimulation. That is, the decrease of inter-hemispheric EEG coherence and the emergence of phasic EMG anticorrelation between arms did not translate into unilateral effects on the dream body. We can only speculate on the rather surprising lack of side-specific effects, and further studies will be important to understand underlying mechanisms. To detect effects on other modalities (e.g. body image distortion, vestibular sensations), a larger group of participants might be necessary. Moreover, the absence of modulatory effects of tDCS on somatosensory experiences, which were reported quite frequently by our participants, could be related to the placement of the tDCS electrodes that was specifically determined by the location of the hand area in the primary motor cortex.

### Implications for consciousness studies

Our study suggests a methodology for identifying, via causal manipulation, the neural correlates of specific types of dream experience. Thus, beyond dream and sleep research, our findings also have more general implications for consciousness research. First, they add another piece of evidence that the neural correlates of specific dream content match the neural correlates of corresponding cognitive and behavioural functions during wakefulness (Siclari et al. 2017). Going beyond mere correlation, our results provide *causal* evidence that the motor cortex is involved in the generation of movement sensations in dreams.

Our results also shed light on the phenomenological profile of self-representation in dreams. In simulation theories, the subjective sense of presence, or the experience of a self in a world, is central to dreaming. While this highlights the importance of self-simulation, the precise pattern of self-experience in dreams, as compared to wakefulness, raises questions (Windt 2015). One possibility is that bodily experience in dreams replicates waking experience; another is that dreams are characterized by a comparative overrepresentation of movement and an underrepresentation of other types of bodily experience (e.g. tactile, thermal, or pain sensations). Our finding that tDCS selectively altered dream movement, taken together with the comparatively low frequency of other types of bodily experience in dreams, is consistent with the second possibility. Future studies could aim to further investigate this question by gathering reports of bodily experience in both dreams and wakefulness, enabling a more direct comparison.

A related question concerns the relation between bodily experiences in dreams and the sleeping physical body. It is commonly thought that dream experience, including bodily experience, is completely independent of outward muscular activity and stimulation of the physical body. However, there are empirical and theoretical reasons for thinking that varying degrees of concordance between dream experience and the physical body exist, on both the levels of sensory input and motor output (Windt et al. 2016; Windt 2018). Lesion studies in cats have shown that pontine lesions, which eliminate REM-sleep related muscular atonia, induce organized motor behavior, such as searching and attacking, during REM sleep (Henley and Morrison 1974; Sastre and Jouvet 1979), possibly indicating dream behaviours. Further examples include (illusory) own-body perception, such as when stimulation to the sleeping body is incorporated in dreams (Nielsen 1993; Sauvageau et al. 1998), and dream enactment behaviors in humans, in which outward muscular activity corresponds to movement sensations in dreams. REM sleep behavior disorder, in which seemingly goal-directed behaviors during REM sleep (such as attacking one’s sleeping partner, attempting to run, etc.) match subjective dream reports, is an extreme example (Schenck et al. 1986; Valli et al. 2012; Howell and Schenck. 2015). But REM sleep is also accompanied by subtler muscular activity in the form of twitching (Blumberg and Plumeau 2016). Its concordance with dream experience seems plausible but has not been systematically investigated.

In our study, bihemispheric tDCS during REM sleep modulated not only dream movement but also outward muscular activity in the arms. Due to the absence of movement reports in several participants, we could not reliably relate individual variance in subjective movement reports to electrophysiological measures. However, our findings are consistent with the possibility that changes in dream movement are related to changes in outward muscular activity during REM sleep. A promising avenue for future research could be to investigate the relevance of bihemispheric tDCS for several movement-related sleep disorders. REM sleep behaviour disorder would be a good place to start because of the match between dream movements and outward physical activity. Other disorders that could benefit from the inhibition of motor activity include sleep walking and restless leg syndrome. Here, however, the association with dream experience is less clear and should be investigated more directly.

### Limitations and outlook

Despite these promising results, the current study has several limitations. First, the effects of tDCS on mental states have been repetitively challenged by replicability difficulties (Tremblay et al. 2014; Horvath et al. 2015a, 2015b) and should thus be treated with caution. Nevertheless, given that motor cortex tDCS during wakefulness provides the most reliable effects (Horvath et al. 2015b; Buch et al. 2017), we expect the same to hold during REM sleep. Second, due to the very complicated and time-intensive protocol of the study, we could only recruit a rather small number of participants. Thus, larger samples and replication studies will be needed in future (Minarik et al. 2016). Furthermore, and again due to the complexity of the setup, we did not include a control stimulation site nor did we switch the side of the bihemispheric stimulation (to left anodal, right cathodal stimulation), which would be especially interesting to disentangle hemisphere-specific effects. Future studies with a larger sample of participants should also explore whether bihemispheric tDCS during REM sleep interferes with a wider range of EEG frequencies involved in motor processing, including alpha and gamma bands as well as broadband responses (Ball et al. 2008; Babiloni et al. 2016).

## Conclusions

To conclude, this study provided, in a controlled setup, evidence that stimulation over the sensorimotor cortex modulates dream content in healthy participants during REM sleep. This has important implications for various research fields, including consciousness research, and sleep and dream research. Future studies will have to pinpoint more specifically which neural mechanisms underlie the inhibition of repetitive movements of the dream self and whether the observed subjective and neurophysiological effects are sufficiently long-lasting to warrant clinical studies in, for example, parasomnia patients.

## Acknowledgements

The study was funded by the Volkswagen Foundation (Project No. I/82 897). Individually, VN was supported by the Signe and Ane Gyllenberg Foundation. JW is the recipient of an Australian Research Council Discovery Early Career Researcher Award. BL is supported by the Swiss National Science Foundation. We thank Jasmine Ho for the language corrections and comments.

## Author Contributions

V.N., J.M.W., A.A.K., T.B. and B.L. conceived the study and designed the experiments. V.N., T.S. and R.P. conducted experiments. V.N., T.B., B.L. analyzed and interpreted EEG data. M.K. and T.B. analyzed and interpreted EMG data. V.N., J.M.W., K.V. and B.L. analyzed and interpreted dream data. V.N., J.M.W., M.K., K.V., T.S., R.P., A.R., T.B., and prepared the manuscript.

## SUPPLEMENTARY MATERIALS

### Content analysis of movement sensations in verbal dream reports: Results

Movements were reported in 49.8% (SEM=10) of dreams following sham-stimulation and 54.9% (SEM=10.9) of dreams following tDCS. Repetitive actions were the most common type of movement, followed by single actions, with passive movements being the least common (see Tables S1 and S2), replicating the pattern observed in the BED Questionnaire data. However, there were no significant differences between the sham-stimulation and tDCS conditions (see Table S2), in contrast to the effects observed in the questionnaire data (see Table 1). The discrepancy could be due to a considerably smaller proportion of explicitly expressed movements in free dream reports compared to the BED Questionnaire answers, i.e. participants tended to omit movements from the spontaneous verbal reports unless asked about them explicitly.

The difference between questionnaire results and dream report analyses has also been found for emotions. The frequency of emotions increases 10-fold if participants are asked to report emotions on a line-by-line basis, as compared to free dream reports (Merritt et al. 1994). When participants are asked to rate the kinds of emotions experienced in their dreams, they specifically report more positive emotions than when their dream reports are analyzed by independent judges (Sikka et al. 2014, 2017). This discrepancy raises important methodological issues that to date have not been fully resolved, and both methods likely have weaknesses and suffer from different kinds of bias (Sikka et al. 2017). One reason for the discrepancy, however, could be that free dream reports lack the focus to allow independent judges to pick up on specific aspects of dream phenomenology, such as emotions or movements. By contrast, when participants’ focus is directed to these aspects, such as through use of questionnaires, this leads to more precise reporting. In our data, similar proportions of different types of movements between external ratings and questionnaire responses, together with the fact that movements are reported more frequently in the questionnaire data, makes us lean towards this interpretation. There are also likely differences in what is reported: in free dream reports, individual movements need to be described in some detail for them to be rated by external judges. By contrast, in the questionnaire, participants rate the occurrence and frequency of specific movement types over the entire dream. Again, this may lead to a more comprehensive picture, but also bears the danger of overgeneralizing.

Nevertheless, the proportion of repetitive actions correlated strongly between the free dream reports and the BED Questionnaire answers in the sham-stimulation condition (Spearman rank order correlation: rho=0.81, p_B-8_=0.033), indicating a strong convergence between these two types of measurement. Interestingly, this association did not hold in the tDCS condition (rho=-0.19, p_B-8_=1). No other correlations were significant.

**Figure S1.**
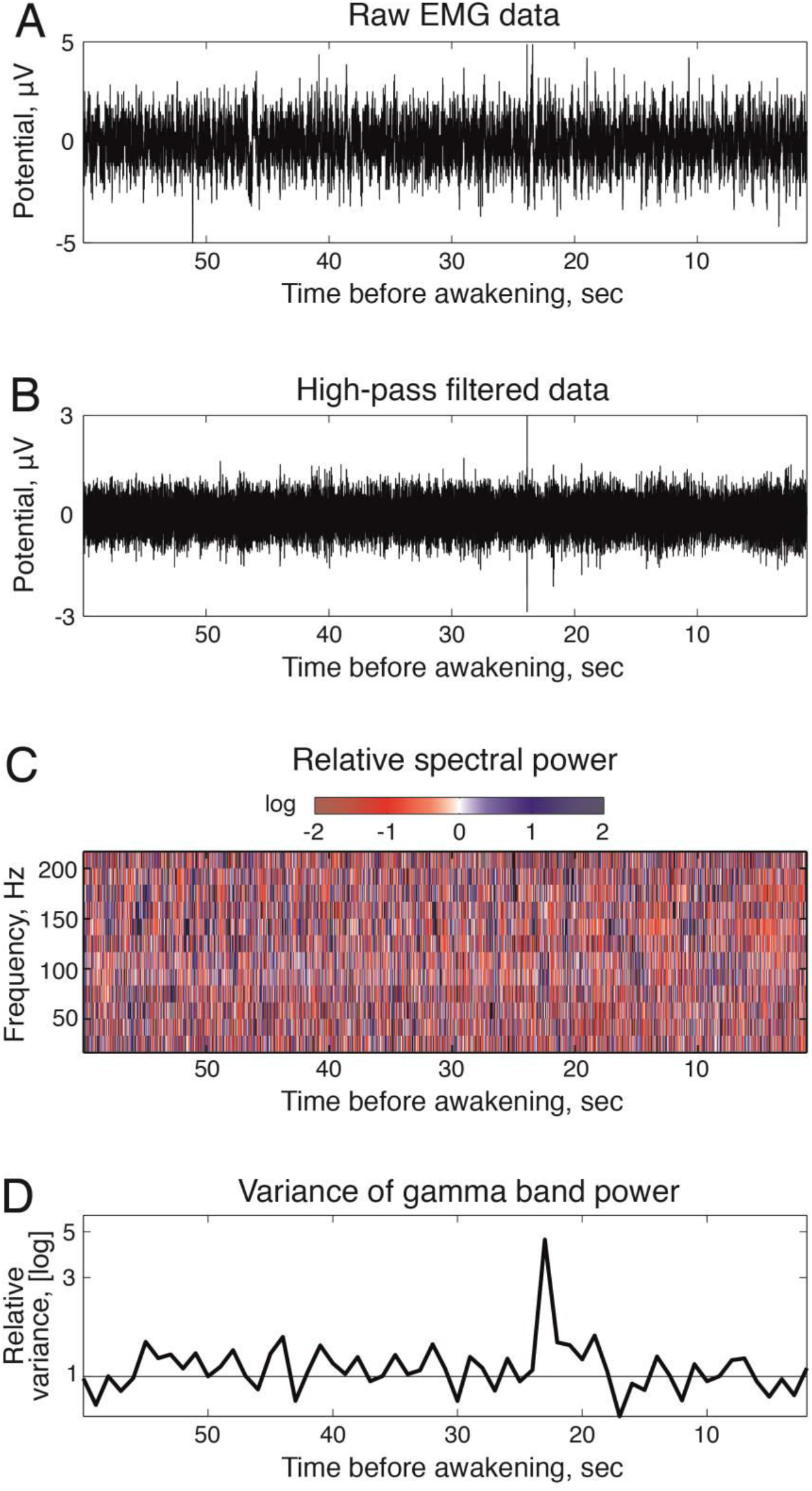
Analysis of peripheral EMG activity. **a)** Exemplary 60 sec EMG recording of the right hand flexoris muscle between termination of tDCS and the awakening. **b)** The same EMG recording after a high-pass filter with a 50 Hz cutoff-frequency. **c)** Relative spectral power of the high-pass filtered EMG recording. **d)** Relative variance of gamma band power, i.e., divided by the average variance of gamma band power in the baseline time window. Values greater than one (above the grey solid line) depict 1 sec segments with a shift towards phasic EMG, values equal or smaller than one depict segments with a shift towards tonic EMG.

**Table S1.** Examples of different types of movement reported in verbal dream reports

| *Movement types* | *Sham* | *tDCS* |
| --- | --- | --- |
| Single actions | “I [...] took away the wires” (P4, N2, A2) | “I was diving” (P1, N1, A1) |
|  | “I was putting together some board” (P10, N2, A2) | “I was [...] to take a pose” (P5, N2, A1) |
|  | “We [...] sat down” (P10, N2, A2) | “I painted a sunset and there was a ship” (P6, N2, A3) |
|  | “I hugged her” (P10, N2, A3) | “I jumped there to the movie” (P7, N1, A1) |
| Repetitive actions | “I remember rubbing quite hard [...] my leg” (P1, N2, A1) | “I was swimming in a pool” (P1, N1, A1) |
|  | “I was walking there” (P3, N2, A2) | “I [...] was writing something” (P3, N1, A1) |
|  | “we are running away from him” (P4, N2, A1) | “we [...] went to the bathroom” (P5, N2, A2) |
|  | “I had been sleepwalking” (P4, N2, A2) | “I was cleaning a table” (P5, N2, A2) |
|  | “I was digging the vegetable garden” (P6, N1, A3) | “I was climbing upstairs” (P7, N2, A2) |
|  | “I was coming out from some room” (P10, N2, A2) | “I was scratching [our cat]” (P8, N2, A3) |
| Passive movements | “we were coming from Lappeenranta with a train” (P4, N2, A1) | “our father was driving me and my brother [...] with a car” (P6, N2, A1) |
|  | “they somehow forced to put my hand to fist” (P7, N1, A1) | “he took my hand and pulled me to the middle” (P6, N2, A1) |

*Note.* P – participant (1-10), N – night (1-2), A – awakening (1-3).

**Table S2.** Dream content analysis: Percentage of dream reports containing movement sensations following sham-stimulation and tDCS during REM sleep

|  | *Sham* | *tDCS* | *Statistical test* |  |
| --- | --- | --- | --- | --- |
|  | *M (SEM)* | *M (SEM)* | *t/Z* | *p* |
| *Movement* |  |  |  |  |
|  | 49.8 (10) | 54.9 (10.9) | t(9) = 0.31 | 0.77 |
| *Movement sub-scales* |  |  |  |  |
| Single actions | 21.6 (10.5) | 24.9 (8.6) | t(9) = 0.19 | 0.85 |
| Repetitive actions | 38.2 (12.7) | 43.2 (9.4) | Z = 0.54 | 0.59 |
| Passive movements | 16.6 (7) | 8.3 (5.7) | Z = 0.76 | 0.45 |

*Note.* t: paired samples t test; Z: Wilcoxon signed-rank test. Uncorrected p values.

**Table S3.** Balance of awakenings between the first and the second night and between the sham-stimulation and tDCS conditions

| *Participant* | *1<sup>st</sup> night, N* | *2<sup>nd</sup> night, N* | *Sham, N* | *tDCS, N* |
| --- | --- | --- | --- | --- |
| 1 | 2 | 3 | 3 | 2 |
| 2 | 2 | 3 | 2 | 3 |
| 3 | 2 | 2 | 2 | 2 |
| 4 | 3 | 2 | 2 | 3 |
| 5 | 3 | 2 | 3 | 2 |
| 6 | 3 | 3 | 3 | 3 |
| 7 | 3 | 3 | 3 | 3 |
| 8 | 3 | 3 | 3 | 3 |
| 9 | 2 | 3 | 3 | 2 |
| 10 | 3 | 2 | 2 | 3 |
| *Total:* | 25 | 25 | 25 | 25 |

## References

Axelrod V, Rees G, Lavidor M, Bar M. 2015. Increasing propensity to mind-wander with transcranial direct current stimulation. Proc Natl Acad Sci. 112:3314–3319.

Babiloni C, Del Percio C, Vecchio F, Sebastiano F, Di Gennaro G, Quarato PP, Morace R, Pavone L, Soricelli A, Noce G, Esposito V, Rossini PM, Gallese V, Mirabella G. 2016. Alpha, beta and gamma electrocorticographic rhythms in somatosensory, motor, premotor and prefrontal cortical areas differ in movement execution and observation in humans. Clin Neurophysiol. 127:641–654.

Baldridge BJ. 1966. Physical concomitants of dreaming and the effect of stimulation on dreamsitle. Ohio State Med J. 62:1272–1275.

Baldridge BJ, Whitman R, Kramer M. 1965. The concurrence of fine muscle activity and rapid eye movements during sleep. Psychosom Med. 27:19–26.

Ball T, Demandt E, Mutschler I, Neitzel E, Mehring C, Vogt K, Aertsen A, Schulze-Bonhage A. 2008. Movement related activity in the high gamma range of the human EEG. Neuroimage. 41:302–310.

Blumberg MS. 2015. Developing Sensorimotor Systems in Our Sleep. Curr Dir Psychol Sci. 24:32–37.

Blumberg MS, Plumeau AM. 2016. A new view of “dream enactment” in REM sleep behavior disorder. Sleep Med Rev. 30:34–42.

Buch ER, Santarnecchi E, Antal A, Born J, Celnik PA, Classen J, Gerloff C, Hallett M, Hummel FC, Nitsche MA, Pascual-Leone A, Paulus WJ, Reis J, Robertson EM, Rothwell JC, Sandrini M, Schambra HM, Wassermann EM, Ziemann U, Cohen LG. 2017. Effects of tDCS on motor learning and memory formation: A consensus and critical position paper. Clin Neurophysiol. 128:589–603.

Buysse DJ, Reynolds CF, Monk TH, Berman SR, Kupfer DJ. 1989. The Pittsburgh sleep quality index: A new instrument for psychiatric practice and research. Psychiatry Res. 28:193–213.

Cicogna P, Bosinelli M. 2001. Consciousness during dreams. Conscious Cogn. 10:26–41.

Cipolli C, Ferrara M, De Gennaro L, Plazzi G. 2017. Beyond the neuropsychology of dreaming: Insights into the neural basis of dreaming with new techniques of sleep recording and analysis. Sleep Med Rev. 35:8–20.

Dang-Vu TT, Desseilles M, Albouy G, Darsaud A, Gais S, Rauchs G, Schabus M, Sterpenich V, Vandewalle G, Schwartz S, Maquet P. 2005. Dreaming: A neuroimaging view. Schweizer Arch fur Neurol und Psychiatr. 156:415–425.

Delorme A, Makeig S. 2004. EEGLAB: An open source toolbox for analysis of single-trial EEG dynamics including independent component analysis. J Neurosci Methods. 134:9–21.

Dement W, Wolpert EA. 1958. The relation of eye movements, body motility, and external stimuli to dream content. J Exp Psychol. 55:543–553.

Dresler M, Koch SP, Wehrle R, Spoormaker VI, Holsboer F, Steiger A, Sämann PG, Obrig H, Czisch M. 2011. Dreamed movement elicits activation in the sensorimotor cortex. Curr Biol. 21:1833–1837.

Fairley JA, Georgoulas G, Mehta NA, Gray AG, Bliwise DL. 2012. Computer detection approaches for the identification of phasic electromyographic (EMG) activity during human sleep. Biomed Signal Process Control. 7:606–615.

Fein G, Raz J, Brown FF, Merrin EL. 1988. Common reference coherence data are confounded by power and phase effects. Electroencephalogr Clin Neurophysiol. 69:581–584.

Feurra, M., Bianco G, Polizzotto NR, Innocenti I, Rossi A, Rossi S. 2011. Cortico-cortical connectivity between right parietal and bilateral primary motor cortices during imagined and observed actions: A combined TMS/tDCS study. Front Neural Circuits. 5:10.

Fortuna M, Teixeira S, Machado S, Velasques B, Bittencourt J, Peressutti C, Budde H, Cagy M, Nardi AE, Piedade R, Ribeiro P, Arias-Carrión O. 2013. Cortical reorganization after hand immobilization: The beta qEEG spectral coherence evidences. PLoS One. 8:e79912.

Fox KCR, Nijeboer S, Solomonova E, Domhoff GW, Christoff K. 2013. Dreaming as mind wandering: evidence from functional neuroimaging and first-person content reports. Front Hum Neurosci. 7:412.

Gandiga PC, Hummel FC, Cohen LG. 2006. Transcranial DC stimulation (tDCS): A tool for double-blind sham-controlled clinical studies in brain stimulation. Clin Neurophysiol. 117:845–850.

Gerloff C, Richard J, Hadley J, Schulman AE, Honda M, Hallett M. 1998. Functional coupling and regional activation of human cortical motor areas during simple, internally paced and externally paced finger movements. Brain. 121:1513–1531.

Henley K, Morrison AR. 1974. A reevaluation of the effects of lesions of the pontine tegmentum and locus coeruleus on phenomena of paradoxical sleep in the cat. Acta Neurobiol Exp (Wars). 34:215–232.

Hobson JA. 1988. The dreaming brain. New York: Basic Books.

Hoff H. 1929. Zusammenhang von Vestibulärfunktion, Schlafstellung und Traumleben. Eur Neurol. 71:366–372.

Hoff H, Pötzl O. 1937. Über die labyrinthären Beziehungen von Flugsensationen und Flugträumen. Eur Neurol. 97:193–211.

Horvath JC, Forte JD, Carter O. 2015a. Quantitative review finds no evidence of cognitive effects in healthy populations from single-session transcranial direct current stimulation (tDCS). Brain Stimul. 8:535–550.

Horvath JC, Forte JD, Carter O. 2015b. Evidence that transcranial direct current stimulation (tDCS) generates little-to-no reliable neurophysiologic effect beyond MEP amplitude modulation in healthy human subjects: A systematic review. Neuropsychologia. 66:213–236.

Howell MJ, Schenck. CH. 2015. REM sleep behavior disorder. In: Videnovic A,, Högl B, editors. Disorders of Sleep and Circadian Rhythms in Parkinson’s Disease. Vienna: Springer. p. 131–144.

Hummel F, Celnik P, Giraux P, Floel A, Wu WH, Gerloff C, Cohen LG. 2005. Effects of non-invasive cortical stimulation on skilled motor function in chronic stroke. Brain. 128:490–499.

Jakobson AJ, Conduit RD, Fitzgerald PB. 2012. Investigation of visual dream reports after transcranial direct current stimulation (tDCS) during REM sleep. Int J Dream Res. 5:87–93.

Jakobson AJ, Fitzgerald PB, Conduit R. 2012a. Induction of visual dream reports after transcranial direct current stimulation (tDCs) during Stage 2 sleep. J Sleep Res. 21:369–379.

Jakobson AJ, Fitzgerald PB, Conduit R. 2012b. Investigation of dream reports after transcranial direct current stimulation (tDCs) during slow wave sleep (SWS). Sleep Biol Rhythms. 10:169–178.

Jasper HH. 1958. The ten-twenty electrode system of the International Federation. Electroencephalogr Clin Neurophysiol. 10:371–375.

Jenkinson N, Brown P. 2011. New insights into the relationship between dopamine, beta oscillations and motor function. Trends Neurosci. 34:611–618.

Khanna P, Carmena JM. 2015. Neural oscillations: Beta band activity across motor networks. Curr Opin Neurobiol. 32:60–67.

LaBerge S, Baird B, Zimbardo PG. 2018. Smooth tracking of visual targets distinguishes lucid REM sleep dreaming and waking perception from imagination. Nat Commun. 9:3298.

Leocani L, Toro C, Manganotti P, Zhuang P, Hallett M. 1997. Event-related coherence and event-related desynchronization/synchronization in the 10 Hz and 20 Hz EEG during self-paced movements. Electroencephalogr Clin Neurophysiol - Evoked Potentials. 104:199–206.

Leslie K, Ogilvie R. 1996. Vestibular dreams: The effect of rocking on dream mentation. Dreaming. 6:1–16.

Lindenberg R, Nachtigall L, Meinzer M, Sieg MM, Floel A. 2013. Differential Effects of Dual and Unihemispheric Motor Cortex Stimulation in Older Adults. J Neurosci. 33:9176–9183.

Lindenberg R, Sieg MM, Meinzer M, Nachtigall L, Flöel A. 2016. Neural correlates of unihemispheric and bihemispheric motor cortex stimulation in healthy young adults. Neuroimage. 140:141–149.

Maquet P, Laureys S, Peigneux P, Fuchs S, Petiau C, Phillips C, Aerts J, Del Fiore G, Degueldre C, Meulemans T, Luxen A, Franck G, Van Der Linden M, Smith C, Cleeremans A. 2000. Experience-dependent changes in changes in cerebral activation during human REM sleep. Nat Neurosci. 3:831–836.

Marshall L. 2004. Transcranial Direct Current Stimulation during Sleep Improves Declarative Memory. J Neurosci. 24:9985–9992.

Matsumoto J, Fujiwara T, Takahashi O, Liu M, Kimura A, Ushiba J. 2010. Modulation of mu rhythm desynchronization during motor imagery by transcranial direct current stimulation. J Neuroeng Rehabil. 7:27.

Mima T, Matsuoka T, Hallett M. 2000. Functional coupling of human right and left cortical motor areas demonstrated with partial coherence analysis. Neurosci Lett. 287:93–96.

Minarik T, Berger B, Althaus L, Bader V, Biebl B, Brotzeller F, Fusban T, Hegemann J, Jesteadt L, Kalweit L, Leitner M, Linke F, Nabielska N, Reiter T, Schmitt D, Spraetz A, Sauseng P. 2016. The importance of sample size for reproducibility of tDCS effects. Front Hum Neurosci. 10:453.

Nam CS, Jeon Y, Kim YJ, Lee I, Park K. 2011. Movement imagery-related lateralization of event-related (de)synchronization (ERD/ERS): Motor-imagery duration effects. Clin Neurophysiol. 122:567–577.

Neuper C, Scherer R, Reiner M, Pfurtscheller G. 2005. Imagery of motor actions: Differential effects of kinesthetic and visual-motor mode of imagery in single-trial EEG. Cogn Brain Res. 25:668–677.

Nielsen TA. 1993. Changes in the Kinesthetic Content of Dreams Following Somatosensory Stimulatipn of Leg Muscles During REM Sleep. Dreaming. 3:99–113.

Nielsen TA. 2017. Microdream neurophenomenology. Neurosci Conscious. 3:nix001.

Nir Y, Tononi G. 2010. Dreaming and the brain: from phenomenology to neurophysiology. Trends Cogn Sci. 14:88–100.

Nitsche MA, Cohen LG, Wassermann EM, Priori A, Lang N, Antal A, Paulus W, Hummel F, Boggio PS, Fregni F, Pascual-Leone A. 2008. Transcranial direct current stimulation: State of the art 2008. Brain Stimul. 1:206–223.

Nitsche M, Paulus W. 2000. Excitability changes induced in the human motor cortex by weak transcranial direct current stimulation. J Physiol. 527:633–639.

Noreika V, Windt JM, Lenggenhager B, Karim AA. 2010. New perspectives for the study of lucid dreaming: From brain stimulation to philosophical theories of self-consciousness. Int J Dream Res. 3:36–45.

Occhionero M, Cicogna P, Natale V, Esposito MJ, Bosinelli M. 2005. Representation of self in SWS and REM dreams. Sleep Hypn. 7:77–83.

Oldfield RC. 1971. The assessment and analysis of handedness: The Edinburgh inventory. Neuropsychologia. 9:97–113.

Opitz A, Yeagle E, Thielscher A, Schroeder C, Mehta AD, Milham MP. 2018. On the importance of precise electrode placement for targeted transcranial electric stimulation. Neuroimage. 181:560–567.

Pellegrino G, Tomasevic L, Tombini M, Assenza G, Bravi M, Sterzi S, Giacobbe V, Zollo L, Guglielmelli E, Cavallo G, Vernieri F, Tecchio F. 2012. Inter-hemispheric coupling changes associate with motor improvements after robotic stroke rehabilitation. Restor Neurol Neurosci. 30:497–510.

Percival D, Walden AT. 2000. Wavelet Methods for Time Series Analysis (Cambridge Series in Statistical and Probabilistic Mathematics). Cambridge: Cambridge University Press.

Pompeiano O. 1967. Sensory inhibition during motor activity in sleep. In: Yahr MD, Purpura DP, editors. Neurophysiological basis of normal and abnormal motor activities. New York: Raven Press. p. 323–375.

Priori A. 2003. Brain polarization in humans: A reappraisal of an old tool for prolonged non-invasive modulation of brain excitability. Clin Neurophysiol. 114:589–595.

Quartarone A, Morgante F, Bagnato S, Rizzo V, Sant’Angelo A, Aiello E, Reggio E, Battaglia F, Messina C, Girlanda P. 2004. Long lasting effects of transcranial direct current stimulation on motor imagery. Neuroreport. 15:1287–1291.

Revonsuo A, Salmivalli C. 1995. A content analysis of bizarre elements in dreams. Dreaming. 5:169–187.

Revonsuo A, Tuominen J, Valli K. 2015. The Avatars in the Machine - Dreaming as a Simulation of Social Reality. In: Metzinger T, Windt JM, editors. Open MIND. Frankfurt am Main: MIND Group. p. 32.

Richter L, Neumann G, Oung S, Schweikard A, Trillenberg P. 2013. Optimal Coil Orientation for Transcranial Magnetic Stimulation. PLoS One. 8:e60358.

Ruohonen J, Karhu J. 2010. Navigated transcranial magnetic stimulation. Neurophysiol Clin Neurophysiol. 40:7–17.

Sanchez-Vives M V., Slater M. 2005. From presence to consciousness through virtual reality. Nat Rev Neurosci. 6:332–339.

Sastre JP, Jouvet M. 1979. Oneiric behavior in cats. Physiol Behav. 9:293–308.

Sauvageau A, Nielsen TA, Montplaisir J. 1998. Effects of somatosensory stimulation on dream content in gymnasts and control participants: Evidence of vestibulomotor adaptation in REM sleep. Dreaming. 8:125–134.

Schenck CH, Bundlie SR, Ettinger MG, Mahowald MW. 1986. Chronic behavioral disorders of human REM sleep: A new category of parasomnia. Sleep. 9:293–308.

Schredl M. 2002. Dream recall frequency and openness to experience: A negative finding. Pers Individ Dif. 33:1285–1289.

Schredl M, Atanasova D, Hörmann K, Maurer JT, Hummel T, Stuck BA. 2009. Information processing during sleep: The effect of olfactory stimuli on dream content and dream emotions. J Sleep Res. 18:285–290.

Schwartz S. 2000. A Historical Loop of One Hundred Years: Similarities between 19th Century and Contemporary Dream Research. Dreaming. 10:55–66.

Schwartz S, Maquet P. 2002. Sleep imaging and the neuropsychological assessment of dreams. Trends Cogn Sci. 6:23–30.

Siclari F, Baird B, Perogamvros L, Bernardi G, LaRocque JJ, Riedner B, Boly M, Postle BR, Tononi G. 2017. The neural correlates of dreaming. Nat Neurosci. 20:872–878.

Sikka P, Feilhauer D, Valli KJ, Revonsuo A. 2017. How you measure is what you get: Differences in self- and external ratings of emotional experiences in home dreams. Am J Psychol. 130:367–384.

Sikka P, Valli K, Virta T, Revonsuo A. 2014. I know how you felt last night, or do I? Self- and external ratings of emotions in REM sleep dreams. Conscious Cogn. 25:51–66.

Speth J, Frenzel C, Voss U. 2013. A differentiating empirical linguistic analysis of dreamer activity in reports of EEG-controlled REM-dreams and hypnagogic hallucinations. Conscious Cogn. 22:1013–1021.

Speth J, Speth C. 2016. Motor imagery in REM sleep is increased by transcranial direct current stimulation of the left motor cortex (C3). Neuropsychologia. 86:57–65.

Strauch I, Meier B. 1996. In search of dreams. Results of experimental dream research. Albany, New York: SUNY Press.

Stumbrys T, Erlacher D, Schredl M. 2013. Testing the involvement of the prefrontal cortex in lucid dreaming: A tDCS study. Conscious Cogn. 22:1214–1222.

Tadel F, Baillet S, Mosher JC, Pantazis D, Leahy RM. 2011. Brainstorm: A user-friendly application for MEG/EEG analysis. Comput Intell Neurosci. 879716:1–13.

Tremblay S, Lepage JF, Latulipe-Loiselle A, Fregni F, Pascual-Leone A, Théoret H. 2014. The uncertain outcome of prefrontal tDCS. Brain Stimul. 7:773–783.

Valli K, Frauscher B, Gschliesser V, Wolf E, Falkenstetter T, Schönwald S V., Ehrmann L, Zangerl A, Marti I, Boesch SM, Revonsuo A, Poewe W, Högl B. 2012. Can observers link dream content to behaviours in rapid eye movement sleep behaviour disorder? A cross-sectional experimental pilot study. J Sleep Res. 21:21–29.

Voss U, Holzmann R, Hobson A, Paulus W, Koppehele-Gossel J, Klimke A, Nitsche MA. 2014. Induction of self awareness in dreams through frontal low current stimulation of gamma activity. Nat Neurosci. 17:810–812.

Windt JM. 2015. Dreaming: A conceptual framework for philosophy of mind and empirical research. Cambridge, MA: MIT Press.

Windt JM. 2018. Predictive brains, dreaming selves, sleeping bodies: how the analysis of dream movement can inform a theory of self- and world-simulation in dreams. Synthese. 195:2577–2625.

Windt JM, Nielsen T, Thompson E. 2016. Does Consciousness Disappear in Dreamless Sleep? Trends Cogn Sci. 20:871–882.

Wu MF. 1993. Sensory processing and sensation during sleep. In: Carskadon MA, editor. Encyclopedia of Sleep and Dreaming. New York: Macmillan. p. 533–535.

Wu T, Hallett M. 2005. A functional MRI study of automatic movements in patients with Parkinson’s disease. Brain. 128:2250–2259.

Yousry TA, Schmid UD, Alkadhi H, Schmidt D, Peraud A, Buettner A, Winkler P. 1997. Localization of the motor hand area to a knob on the precentral gyrus. A new landmark. Brain. 120:141–157.

Zaepffel M, Trachel R, Kilavik BE, Brochier T. 2013. Modulations of EEG Beta Power during Planning and Execution of Grasping Movements. PLoS One. 8:e60060.

